# Revealing the nanoscale morphology of the primary cilium using super-resolution fluorescence microscopy

**DOI:** 10.1101/437640

**Authors:** Joshua Yoon, Colin J. Comerci, Lucien E. Weiss, Ljiljana Milenkovic, Tim Stearns, W. E. Moerner

## Abstract

Super-resolution (SR) microscopy has been used to observe structural details beyond the diffraction limit of ~250 *nm* in a variety of biological and materials systems. By combining this imaging technique with both computer-vision algorithms and topological methods, we reveal and quantify the nanoscale morphology of the primary cilium, a tiny tubular cellular structure (~2-6 *μm* long and 200-300 *nm* diameter). The cilium in mammalian cells protrudes out of the plasma membrane and is important in many signaling processes related to cellular differentiation and disease. After tagging individual ciliary transmembrane proteins, specifically Smoothened (SMO), with single fluorescent labels in fixed cells, we use three-dimensional (3D) single-molecule SR microscopy to determine their positions with a precision of 10-25 *nm*. We gain a dense, pointillistic reconstruction of the surfaces of many cilia, revealing large heterogeneity in membrane shape. A Poisson surface reconstruction (PSR) algorithm generates a fine surface mesh, allowing us to characterize the presence of deformations by quantifying the surface curvature. Upon impairment of intracellular cargo transport machinery by genetic knockout or small-molecule treatment of cells, our quantitative curvature analysis shows significant morphological differences not visible by conventional fluorescence microscopy techniques. Furthermore, using a complementary SR technique, 2-color, 2D STimulated Emission Depletion (STED) microscopy, we find that the cytoskeleton in the cilium, the axoneme, also exhibits abnormal morphology in the mutant cells, similar to our 3D results on the SMO-measured ciliary surface. Our work combines 3D SR microscopy and computational tools to quantitatively characterize morphological changes of the primary cilium under different treatments and uses STED to discover correlated changes in the underlying structure. This approach can be useful for studying other biological or nanoscale structures of interest.

## Introduction

Revealing the three-dimensional (3D) nanoscale membrane structure of biological cells in both a non-invasive and precise manner remains a challenging problem. Structural components that define the cell outer morphology have been examined using various conventional fluorescence microscopy methods (1, 2), but the resolution is ultimately limited by diffraction (~250 *nm*). Super-resolution (SR) fluorescence microscopy circumvents the diffraction limit by either imaging and localizing a sparse set of single molecules (SM) separated in time (3–5) or by shrinking the effective excitation point-spread function (PSF) *via* STimulated Emission Depletion (STED) microscopy (6–8). The relatively non-invasive nature of optical microscopy can then be utilized for nanoscale structural analysis.

The first technique, SM super-resolution (SR) microscopy, requires the structure of interest to be densely labeled with a fluorophore that has at least two states with distinct emissive (on-off) properties, and this mechanism is crucial to force sparsity in each imaging frame (9). Active control of the labels can be achieved either optically (*e.g.* illumination with a near-UV light source for photoactivation) (10–13) or non-optically (*e.g.* label reacting with a nearby ligand in a reversible fashion to induce blinking) (14–16). Each molecule is then fit with a mathematical function that determines its position with a precision that scales as 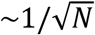 where *N* is the number of photons detected, and the locations are eventually merged together to create a reconstructed image with an enhanced resolution of typically ~20-40 *nm* for fluorescent protein labels, with improved precisions for small-molecule emitters providing more photons. However, determining 3D position information is difficult due to the standard PSF being symmetric about the focal plane and only observable over a relatively small axial range (~500 *nm*) compared to typical structures of interest (17). To address these issues, recent developments in PSF engineering have made it possible to encode depth information within the PSF shape of the SM emitter, including astigmatism (18), multiplane (19), 4Pi (20), and Fourier domain manipulation (21–26). In particular, the double-helix (DH) PSF allows for 3D SM localization, because individual emitters appear as two spots on the camera which revolve around one another in different *z*-positions, thus the angle between the two lobes encodes *z* information over a relatively large axial depth-of-field (~2 *μm*) (27). The DH-PSF is generated by moving the camera away from the intermediate image plane (IIP) by four focal lengths *f*, and then placing an optical phase mask in the collection path equidistant between two lenses of focal length *f*, referred to as a *4f* system (28). This technique has been used to study nanohole arrays across large fields of view (29), protein localization patterns within bacteria (30–32), and organelles inside mammalian cells (33).

An alternate SR microscopy method, STED microscopy (6–8), requires two laser beams which work in concert to shrink the effective PSF of the excitation spot in a confocal scanning configuration. The excitation beam is focused to a diffraction-limited spot onto a sample and pumps the molecules into the first electronic excited state. A second overlapping depletion beam at longer wavelengths is shaped optically into a donut with a dark center, selectively forcing emitters that lie at the edge of the excitation spot to produce far-red stimulated emission light that is filtered out. In this way, only the fluorophores at the very center of the donut undergo normal fluorescence emission, which yields imaging resolution better than the diffraction limit. Images are generated by scanning over the co-aligned laser beams over the region of interest (34, 35). This technique has been used to examine nitrogen-vacancy centres (36), neurons (37), centriole proteins (35, 38), and many other structures at high resolution.

In this work, we employed both imaging techniques to study the primary cilium in cells, which is a tubular structure ~2-6 *μm* long and ~200-300 *nm* in diameter. The overall structure is based on the axoneme (39), a column of nine-fold symmetric microtubule doublets, running through the center of the cilium. The cilium is covered by the ciliary membrane, an extension of the plasma membrane which has a distinct protein-lipid composition to the rest of the cell (40). This non-motile, antenna-like sensory organelle is essential for facilitating proper signal transduction, cell-to-cell communication, and proper regulation of cell division in mammalian cells (41, 42). Impaired ciliary function leads to a collection of human diseases termed “ciliopathies” with a broad range of pathologies of various severities including polydactyly (multiple digits), cystic kidney disease, obesity, and retinal degeneration (43). One reason these debilitating diseases arise is due to impaired expression of intraflagellar transport (IFT) complexes, which is a family of proteins that move components required for constructing the cilium up and down along the structure (44). When IFT is functioning incorrectly, cilia are often found completely missing or misshapen when observed using fluorescence or electron microscopy (EM) (45–48). Although recent studies using SR microscopy have elucidated the distribution of a plethora of proteins residing within the primary cilium (35, 49–51), there remains a need for quantitative approaches to characterize the ciliary membrane shape.

Here we describe a method for revealing and quantifying the membrane structure of the primary cilium using a combination of SR microscopy techniques and computer-vision algorithms. In mouse embryonic fibroblast (MEF) cells, we genetically express and label a crucial transmembrane ciliary protein Smoothened (SMO) with a bright fluorophore, add a blinking buffer for STORM (52), and perform 3D SM SR using the DH-PSF with a precision of 10-25 *nm*. Our processed 3D molecular positions of SMO are then fed into the Poisson Surface Reconstruction (PSR) algorithm, which produces 3D triangulated surface meshes representing the shape of each primary cilium membrane. Computing the mean (*H*) and Gaussian (*K*) curvature provides a quantitative picture of the surface shape at a high resolution. Upon impairment of the retrograde transport machinery through genetic modification or treatment, the narrowing and bulging near the the tip of the cilium is more severe. Surface information allows us to calculate the Willmore Energy (*W_E_*) (53) as an appropriate metric for measuring its overall characteristic shape, and shows a significant increase in our mutant cells. Furthermore, when imaging primary cilia in *IFT25* mutant cells using 2-color 2D STED microscopy, we observe fluorescently labeled αTubulin spanning the entire primary cilium and in some cases occupying the bulging ciliary membrane at the tip, features in which we did not find in wild type MEF cells. By combining SR fluorescence microscopy and quantitative surface meshing, this method of shape analysis may be used to study a broad range of nanoscale tubular structures in a more quantitative and precise way.

## Materials and Methods

### Cell culture

Mouse embryonic fibroblast (MEF) cells were generated, which stably express Smoothened (SMO) proteins with an extracellular SNAP tag (SNAP-SMO) and Pericentrin-YFP (PACT-YFP) as a basal marker. Cell lines used: (1) SMO−/−, SNAP-SMO, YFP-PACT; (2) IFT25−/− (*IFT25*), SNAP-SMO, YFP-PACT as described in Ref. (54).

### Sample preparation (3D SR Microscopy)

MEF cells are first labeled with Benzylguanine-Alexa647 (NEB, S9136S), then fixed with 4% paraformaldehyde (PFA) and finally treated with a quenching solution of 10 *mM* NH_4_Cl. The samples are washed 3x with 1x PBS, pH 7.4 then either imaged immediately or stored at 4°C up to 1-week before being discarded.

### 3D SM Microscopy Imaging

Experiments are performed on a customized inverted microscope (Olympus IX71) where our sample is mounted on a piezo-electric stage (PI-Nano) and is in contact with an oil-immersion objective (Olympus, 100x, 1.4 NA, UPLANSAPO). New imaging buffer is added for each primary cilium imaged (1-2 hours), which consists of glucose oxidase (Sigma-Aldrich, G2133), catalase from bovine serum (Sigma-Aldrich C100), 100 mM Tris-HCl, pH 8.0 (ThermoFisher Scientific, 15568025), 10% (w/v) glucose solution (Sigma-Aldrich, 49139), 140 mM beta-mercaptoethanol (Sigma-Aldrich, M6250), and H_2_O (Nanopure) (52). We locate one primary cilium and image SNAP-SMO-Alexa647 and PACT-YFP using the 641 *nm* (Coherent Cube, 100 *mW*) and 514 *nm* (Coherent Sapphire, 50 *mW*) lasers, respectively. When we place the cilium at the center of the field-of-view (FOV), the double-helix (DH) phase mask is carefully placed at the Fourier Plane (FP) in our *4f* system (Fig. S1 in the Supporting Material). The DH-phase mask has the effect of optically splitting the standard PSF into two lobes, where the midpoint determines its *x, y* position while the angular orientation provides *z* information which is separately calibrated (see Supporting Material). To begin imaging, the intensity of the 641 *nm* laser (1-5 *kW/cm*^2^) is increased, and the collection of labels is allowed to bleach down until single-molecule concentrations are achieved and imaging frames are recorded in the presence of blinking. Over the next hour, irradiation from a secondary 405 *nm* (Obis, 100 *mW*) laser is used to maintain photoblinking labels at suitable densities. Red fluorescent fiducial beads are also imaged simultaneously several microns away from the cilium to correct for sample drift. Detected fluorescence is recorded using a silicon EMCCD camera (Andor Xion, DU-897U-CS0-#BV) at a speed of 20 *frames/second* (50 *ms*/*frame*) with an electron-multiplying gain of 200, and further analyzed using the *easyDHPSF* program, which is freely available online (55).

### Meshing

Using MeshLab (ISTI-CNR, Pisa, Italy), we apply the Poisson Surface Reconstruction algorithm to our 3D data in order to create a triangulated surface as described in the SI. The results are exported in a ‘*.ply’ file format which is used to plot our 3D mesh in MATLAB (The Mathworks, Natick, MA) and for our curvature analysis.

### Curvature Analysis

We then fit a surface to a “patch” of points, which consists of the 1^st^ & 2^nd^ nearest neightbors to each vertex of the mesh (Fig. S4 in the Supporting Material), to the following form:

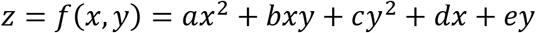

where its coefficients are used to calculate the Mean Curvature (*H*) and Gaussian Curvature (*K*) as follows (56):

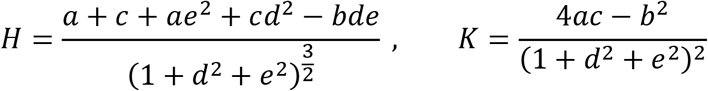

We can then calculate the Willmore Energy (53) by the following expression:

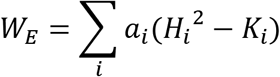

where *H_i_*, *K_i_*, and *a_i_* are the mean curvature, Gaussian curvature, and surface area of the *i^th^* triangle of the mesh respectively. The Willmore Energy Density is defined as *W_E, D_* = *W_E_ /A*, where *A* = ∑_*i*_*a_i_*.

### Sample Preparation for 2D STED

Cell samples are fixed with 4% PFA for 15 *min* at 25°C, washed with 1x PBS, then immersed in a blocking solution, consisting of 1% TritonX-100, Normal Donkey Serum (Jackson ImmunoResearch, 017-000-121), and 1x PBS, for 30 *min* at 25°C. We stain our samples with primary/secondary antibody, targeting SNAP-SMO with Atto647N and αTubulin with Star520SXP.

### 2-color 2D STED Microscopy

STED images were collected on a bespoke 2-color fast scanning STED microscope (Fig. S6 in the Supporting Material). Briefly, laser pulses of 750 *nm* for the depletion beam and 530 *nm* and 635 *nm* for the excitation beams are scanned along the fast axis using a 7.5 *kHz* resonant mirror and along the slow axis using a piezo stage. Fluorescence is detected through a ~0.7 and 0.8 A.U. pinhole (in the red and green channels, respectively) sequentially on a Si APD between 550-615 *nm* for the green channel and 660-705 *nm* for the red channel. The images are acquired using a custom LabVIEW algorithm running on an FPGA. See Supporting Material for details.

## Results and Discussion

### 3D SR Microscopy reveals heterogeneity in membrane shape

To study the ciliary membrane, we chose the transmembrane protein SMO as the labeling target because these proteins are known to move in a largely diffusive manner and be relatively dense during Hh pathway activation (54, 57, 58). Our baseline MEF cells, denoted as *wt*, were treated with a drug named Smoothened Agonist (SAG) (59), which activates the Hh signaling pathway and triggers SMO to localize specifically to the primary cilium (Fig. 1a). SMO is conveniently labeled extracellularly using cells expressing a genetic fusion to the SNAPtag (SNAP), which reacts with a benzyl-guanine-derivatized fluorophore, here Alexa647. After SNAP-SMO is covalently labeled with the Alexa647 fluorophore and chemically fixed, we first imaged our samples, in the presence of a blinking buffer, using a 641 *nm* laser to detect a diffraction-limited image of SNAP-SMO, then used a 514 *nm* laser to detect PACT-YFP, which is a fluorescent marker of the primary cilium base (Fig. 1b). In order to gain as much information of the ciliary surface, a longer-than-normal time (~60 *min*) is spent imaging blinking labels for one sample, where each detected molecule appears as two spots that both roughly take on the shape of a Gaussian on our camera. We capture molecules within a 2 *μm*-thick axial range with a high spatial precision and a relatively large x-y area (40 *μm* x 40 *μm*) which allows us to simultaneously image SNAP-SMO and a nearby bright fiducial marker used to correct for sample drift (Fig. 1c). The single emitters produced a signal of ~8700 photons on average and a mean background level of ~216 photons/pixel, yielding a ~10-25 *nm* spatial resolution with several thousands of localizations for each cilium (Fig. S2 in the Supporting Material). Our complete 3D reconstructions, or “point clouds”, revealed localizations densely-packed within the ciliary membrane in a near homogenous fashion (Fig. 1d). By looking at cilia in 3D on many *wt* MEF cells, we observe a variety of distinct morphological classes, such as bending, kinking, bulging, and the occasional budding with a diameter of ~100 *nm* (Fig. 1c, e, f). In addition, we were also able to detect a few SNAP-SMO molecules outside of the primary cilium, likely on the plasma membrane and were not analyzed further. Despite the fact that these cells were grown under identical conditions, the wide range of heterogeneity in shape from one cilium to another is clearly apparent. Furthermore, our results demonstrate that the underlying structure of the primary cilium is not a simple cylinder/hemispheric-cap as it is frequently modeled.

**Figure 1:**
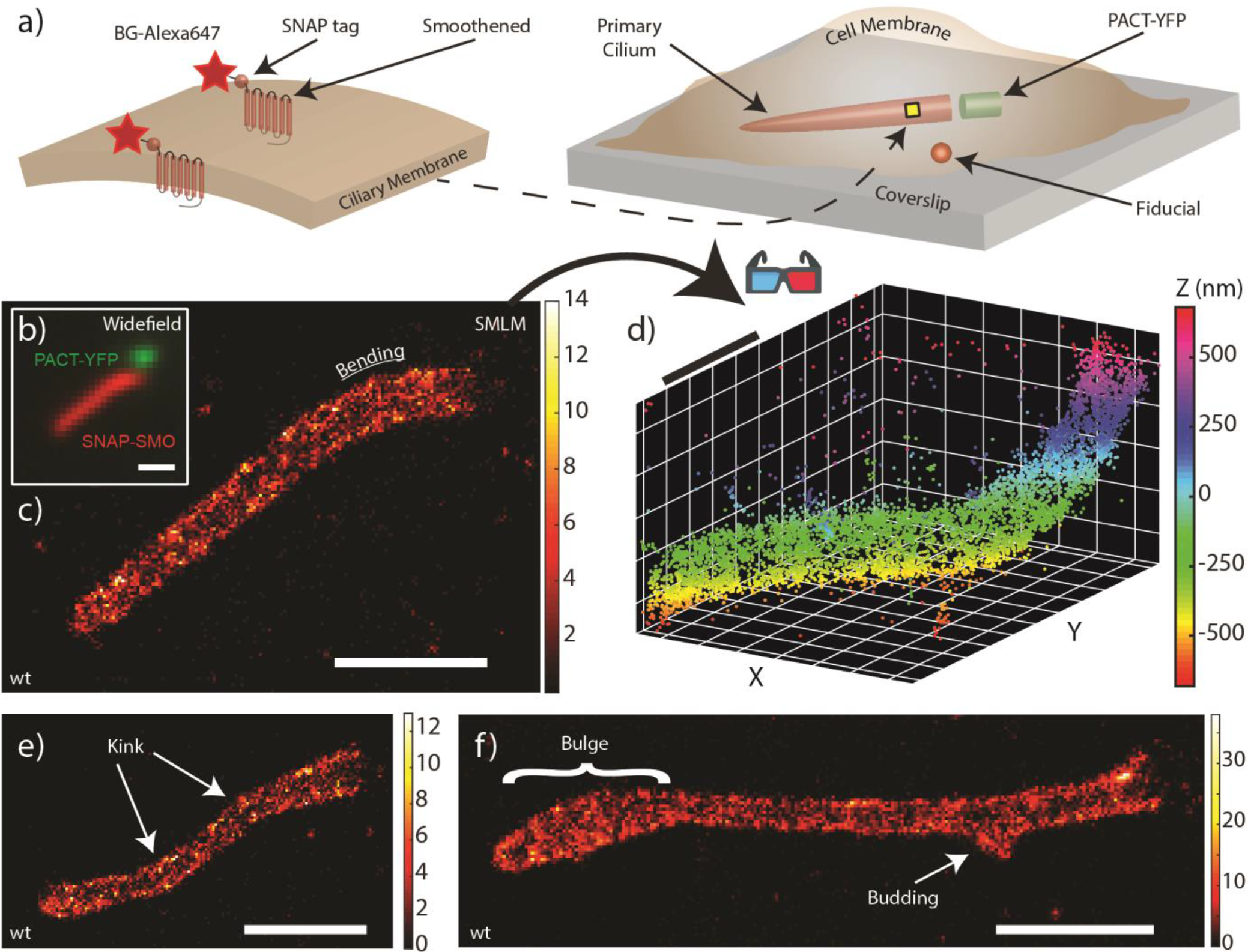
Labeling and 3D SR imaging of the ciliary membrane. (a) SNAP-SMO proteins, where the SNAP protein is on the extracellular side, are covalently labeled with BG-Alexa647 along the ciliary membrane, which are usually found near the coverslip surface. PACT-YFP indicates the base of the cilium and a nearby bright fiducial is used to correct for spatial drift. (b) Overlaid diffraction-limited images of the SNAP-SMO and PACT-YFP in chemically fixed control MEF cells that were treated with SAG. 3D SR microscopy using the double-helix point spread function (DH-PSF) was performed to obtain a localization map of SNAP-SMO molecules along one primary cilium, reconstructed as a (c) 2D histogram and (d) 3D scatterplot. For SNAP-SMO distributions of other primary cilia, there is evidence of (e) kinking, (f) bulging, and budding within the same control MEF cells. Scale bar = 1 μm.

### 2D surface fitting captures the ciliary membrane surface and enables curvature quantification

In order to extract the complex surface of the ciliary membrane, we used the Poisson Surface Reconstruction (PSR) algorithm in the freely available package MeshLab (60). The advantages of this algorithm are that it is robust to noisy, non-uniform point cloud data, and outperforms other commonly used surface fitting methods (61). We prepared our dataset by first selecting only the points that reside within the primary cilium. Because the algorithm works best when the underlying surface is topologically closed, we selected several points near the base of the cilium to determine a plane of best fit. The point cloud was then copied and rotated 180° about the plane axis, producing a rotationally symmetric point cloud made up of two identical shapes. Using the proper fitting parameters, we then generated 2D triangular meshes of each primary cilium, providing us with a clearer picture of the surface (Fig. 2a). Rather than having to navigate through a point cloud, we are able to observe the contours of the surface with greater clarity from many different perspectives (Movie S1, S2 in the Supporting Material). Similar to our super-resolution reconstructed images, we can better identify regions of bulging, narrowing, and enlargement along the ciliary membrane, which are prominent features in *IFT25* mutant cells (Fig. 2b). In addition, the vertices of the mesh allowed us to obtain the ciliary axis, a centroid line that spans from the base to the tip of the primary cilium (see Fig. 3 below). If necessary, surface meshes for wt and mutant cells can also be 3D-printed using standard plastic-filament material, a true 3D representation of the meshes shown in Fig. 2 (Fig. S3 in the Supporting Material).

**Figure 2:**
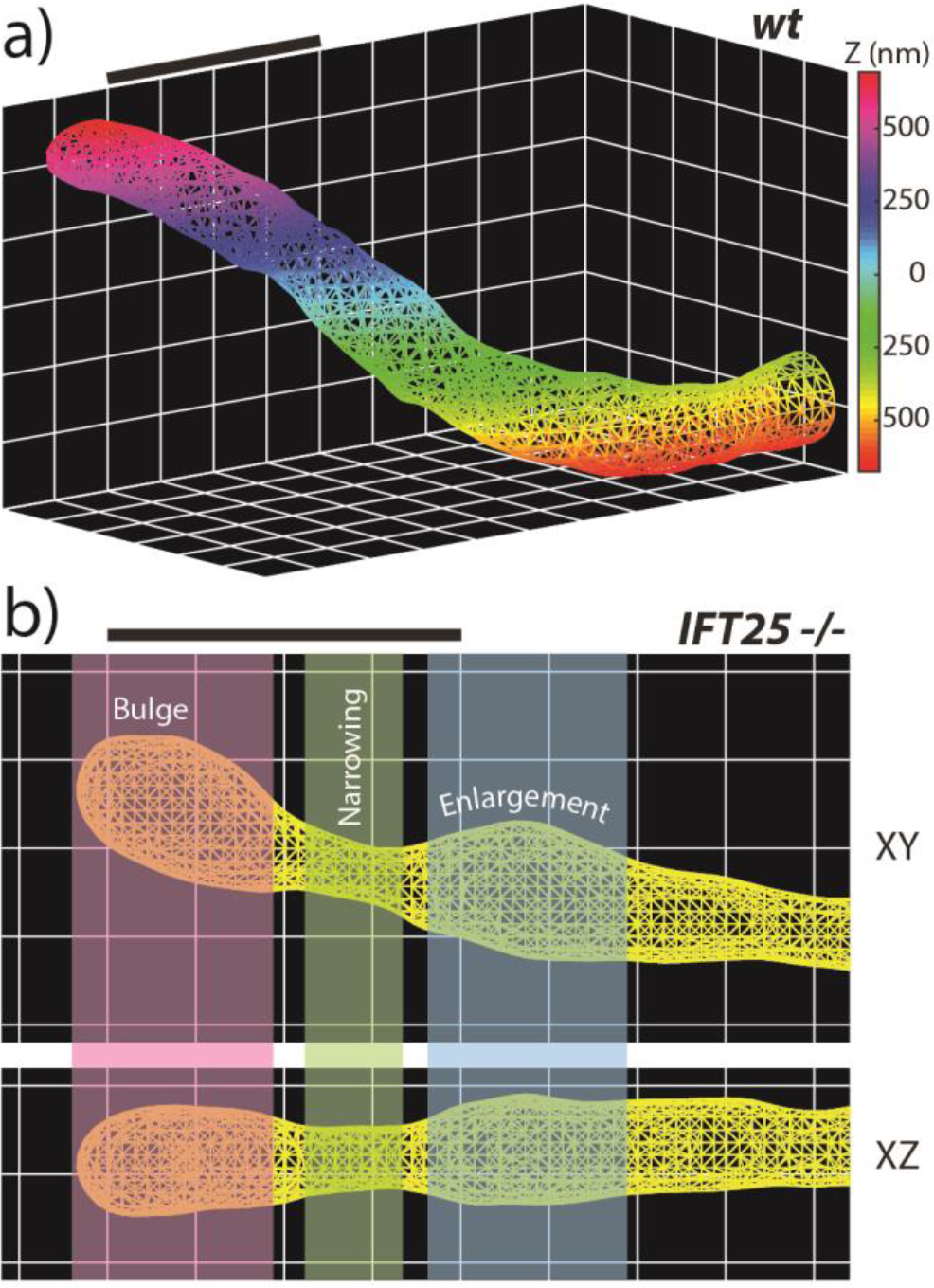
2D surfaces of the ciliary membrane. MeshLab is used to create a triangular mesh using the input point cloud for (a) control MEF primary cilium and (**b**) *IFT25* mutant cell, where different colored regions indicate, from left to right, bulging at the tip of the cilium, narrowing, and then enlargement. Scale bar = 1 μm.

**Figure 3.**
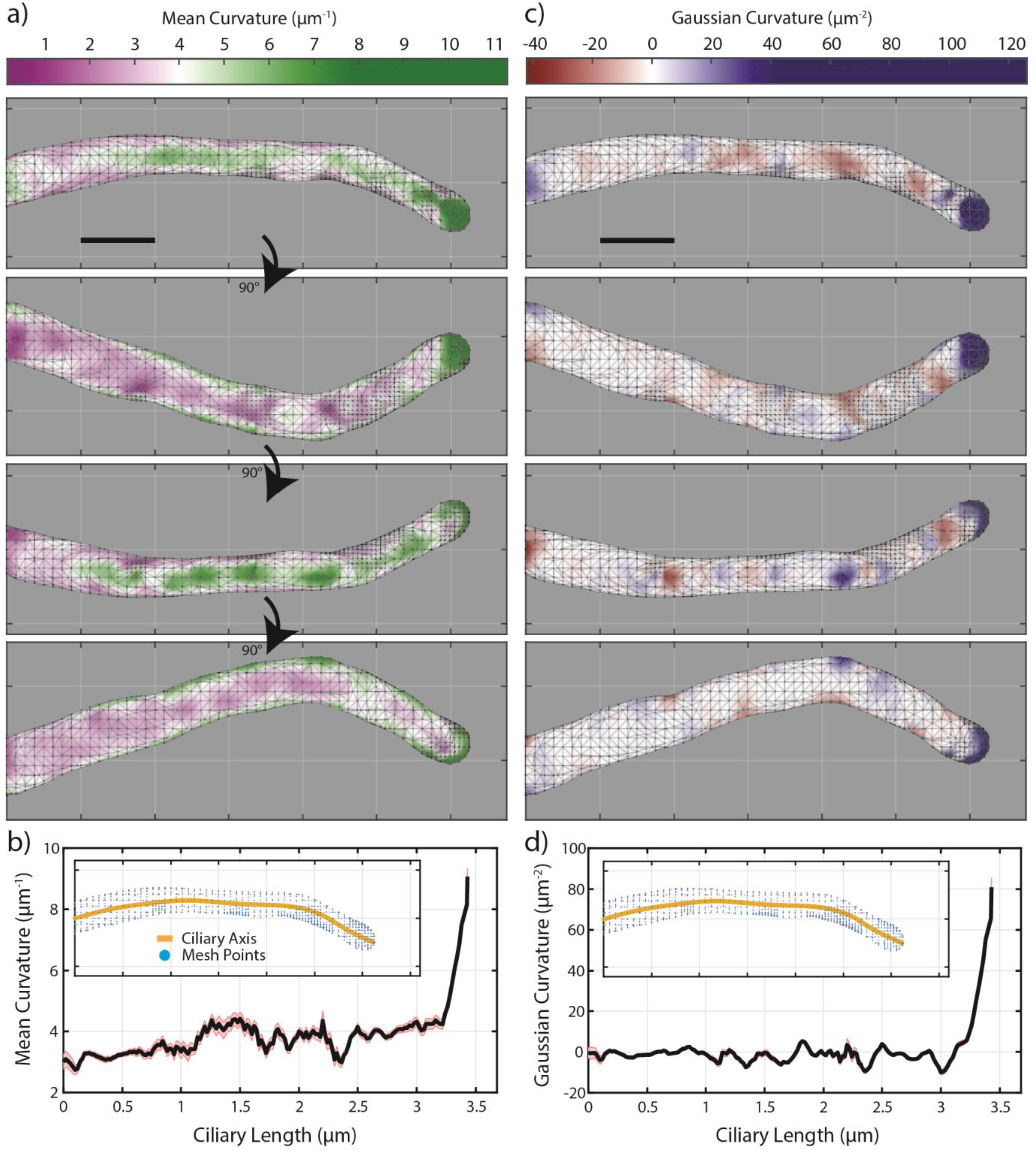
Calculating Mean and Gaussian Curvature along mesh. (a) Mean curvature (*H*) and (c) Gaussian curvature (*K*) are calculated for each vertex along the mesh for a control MEF cell, and displayed here as heat maps. Each panel is a 90° rotation about the horizontal axis. The average (b) *H* and (d) *K* is calculated over a 100-nm wide window that slides along the ciliary axis in 20-nm steps. (Inset figures) Yellow lines indicate the measured ciliary axis; blue points are the vertices of the mesh. Reporting mean ± S.E.M. Scale bar = 0.5 μm.

We then quantified the local shape of the ciliary membrane by calculating its mean (*H*) and Gaussian (*K*) curvature along the entire surface (Fig. 3). A *K* surface heat map highlights regions of curvature where they represent one of the following shapes: (1) sphere, (2) cylinder, or (3) saddle point. When comparing one cilium to another, it is easy to recognize morphological differences and locate where regions of unusual or interesting behavior occur. Further, by measuring the surface area, *A*, and the length of the cilium, *l*, we can then compute an approximate diameter along the shaft of the primary cilium (Table 1). When plotting the values of *A* and *l* cell by cell in a 2D scatterplot, overall they follow a linear relationship, as expected (Fig. S5 in the Supporting Material). None of the diameters are significantly different across all four conditions.

**Table 1:** Measuring the length, surface area, and diameter of the primary cilium for different conditions from 3D SR imaging. Ciliary length is simply the complete length from the tip to the base of the cilium, summing over the areas of all the triangles is the surface area, and the diameter displayed is computed from a model assuming the cilium is a cylinder with a hemispheric cap (Fig. S5, Supporting Material). Reporting mean ± S.T.D.

| | $wt_{SAG}$ | $wt_{SAG}^{cb}$ | $IFT25_{NoAg}$ | $IFT25_{SAG}$ |
| --- | --- | --- | --- | --- |
| Number of Cells | 17 | 14 | 16 | 12 |
| Ciliary Length ($\mu\text{m}$) | $2.821 \pm 0.658$ | $2.564 \pm 0.779$ | $3.170 \pm 0.640$ | $3.083 \pm 0.371$ |
| Surface Area ($\mu\text{m}^2$) | $2.403 \pm 0.631$ | $1.960 \pm 0.610$ | $2.594 \pm 0.728$ | $2.475 \pm 0.349$ |
| Diameter (nm) | $272 \pm 40$ | $244 \pm 29$ | $258 \pm 41$ | $257 \pm 31$ |

To further quantify the cilium’s overall morphology, we calculated its Willmore energy (*W_E_*) which is expressed by the following

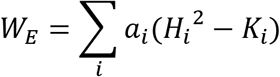

where *H_i_*, *K_i_*, and *a_i_* are the mean curvature, Gaussian curvature, and surface area of the *i^th^* triangle of the mesh respectively. The intuition behind *W_E_* is that it is a strictly positive quantity with a minimum value of 0 when the object is a sphere. For a surface that is characterized to be morphologically more complex, which includes being stretched in a certain direction or severely indented, *W_E_* will take on a larger value. However, a normal cilium that is unusually long with a large surface area can have a value of *W_E_* comparable to one that is relatively short but very narrow and bulbous. To offset the variability in surface area and to ensure a fairer comparison, we computed the total surface area of one mesh, *A* = ∑_*i*_ *a_i_*, and define here the Willmore energy density *W_E, D_* = *W_E_ /A*. By calculating this value for all of our primary cilia, we found it to be a useful metric for comparing overall shape phenotype.

### Altered morphology is detected in cells with impaired retrograde transport

We examined cells where *IFT25* were genetically knocked out, which allows SMO to densely accumulate to the primary cilium despite the absence of pathway activation. 3D SM microscopy on many different cilia in untreated *IFT25* cells shows an altered morphology (Fig. 4c). In particular, there was a larger proportion of cells that exhibited a bulbous tip which was apparent upon generating the surface meshes. We characterized the surfaces by calculating their respective *H* and *K* profiles, where large negative values of *K* correspond to regions of severe narrowing of the mesh (Fig. 4). As a related condition, *IFT25* cells treated with SAG should have an added accumulation of SMO, and these cells showed a similar phenotype to the *IFT25* cells without SAG. This suggests that increased bulging effects in the primary cilium are largely due to the absence of transport complexes and the accumulation of non-SMO proteins, rather than driven strictly by SMO protein accumulation.

**Figure 4:**
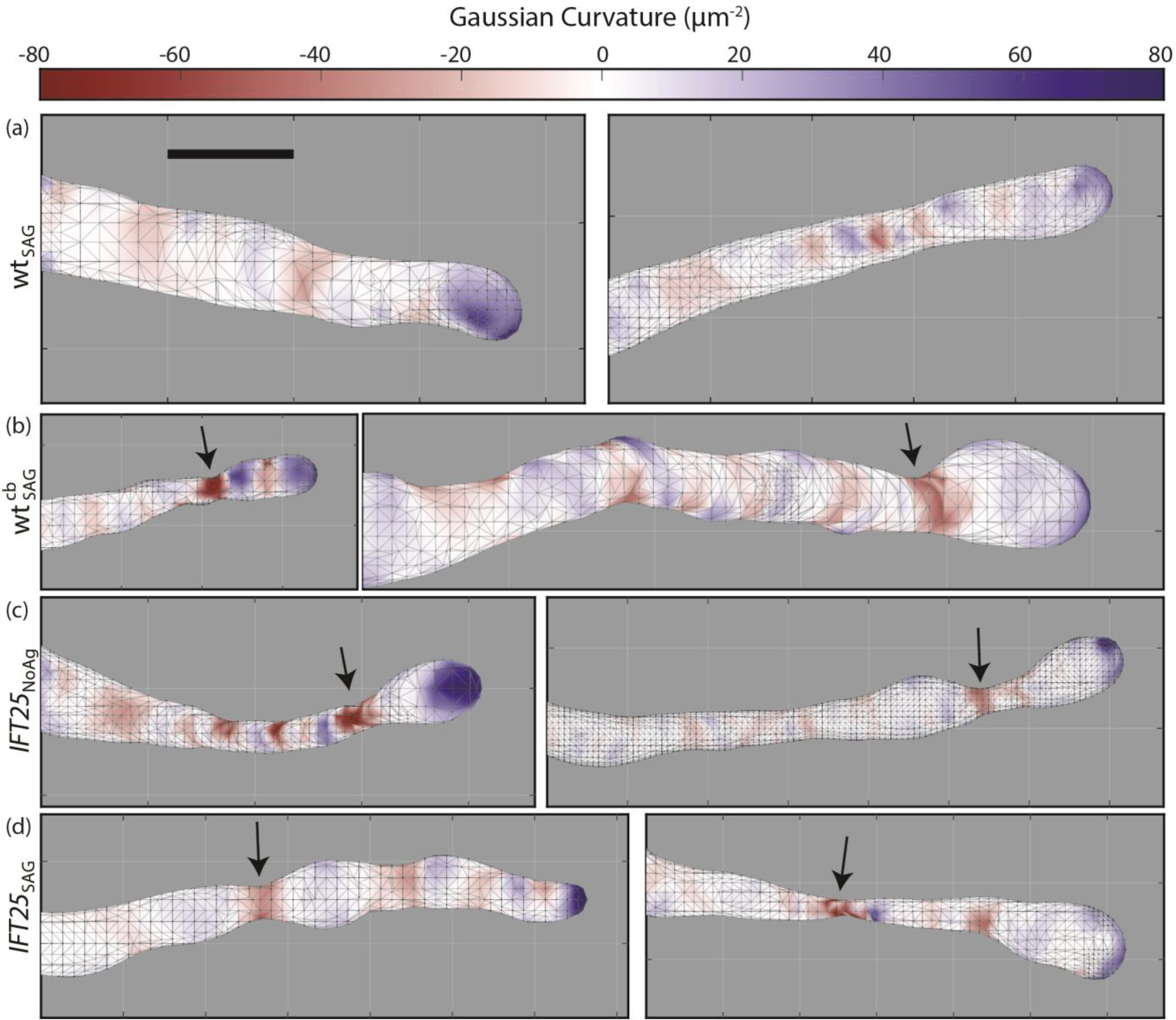
Comparing Gaussian Curvature (*K*) heat maps for different representative primary cilia across all conditions. Largely normal cilia are found among the control MEF cells, while in the mutant cells, there is severe narrowing occurring along the ciliary surface, indicated by the corresponding black arrows and red areas. Heat maps are all on the same scale. Scale bar = 0.5 μm.

To see if we could observe a similar phenotype in our *wt* MEF cells when transport is modified, we treated them with ciliobrevin D and performed 3D SM microscopy. This small-molecule inhibitor of the motor cytoplasmic dynein perturbs retrograde protein trafficking within the primary cilium (62). It has been previously shown that prolonged incubation of cells with ciliobrevin D leads to complete cilia loss, but within a 4-hour incubation period, cilia are still present, although with disrupted retrograde transport (62). As expected (Fig. 4b), in cells treated with ciliobrevin D, increased bulging and narrowing was observed, similar to the morphological features found in *IFT25* mutant cells.

To quantitatively distinguish distinct categories of morphology, we calculated the curvatures and Willmore energies adjacent to a constriction. The most significant contstriction occurs at the position of minimum Gaussian curvature *K* (*K_min_*). By computing the average *K*, (*K̅*) at the constriction and on both sides of the position of the *K_min_* value, the average curvature is significantly smaller for each of the three mutant cases compared to the *wt* condition, indicating a more severe narrowing near this region (Fig. 5a). Furthermore, when calculating *W_E, D_* for every cilium under the different mutant conditions, the value was on average higher among *IFT25* cells compared to the *wt* case, and was also higher when *wt* cells are treated with ciliobrevin D (Fig. 5b).

**Figure 5:**
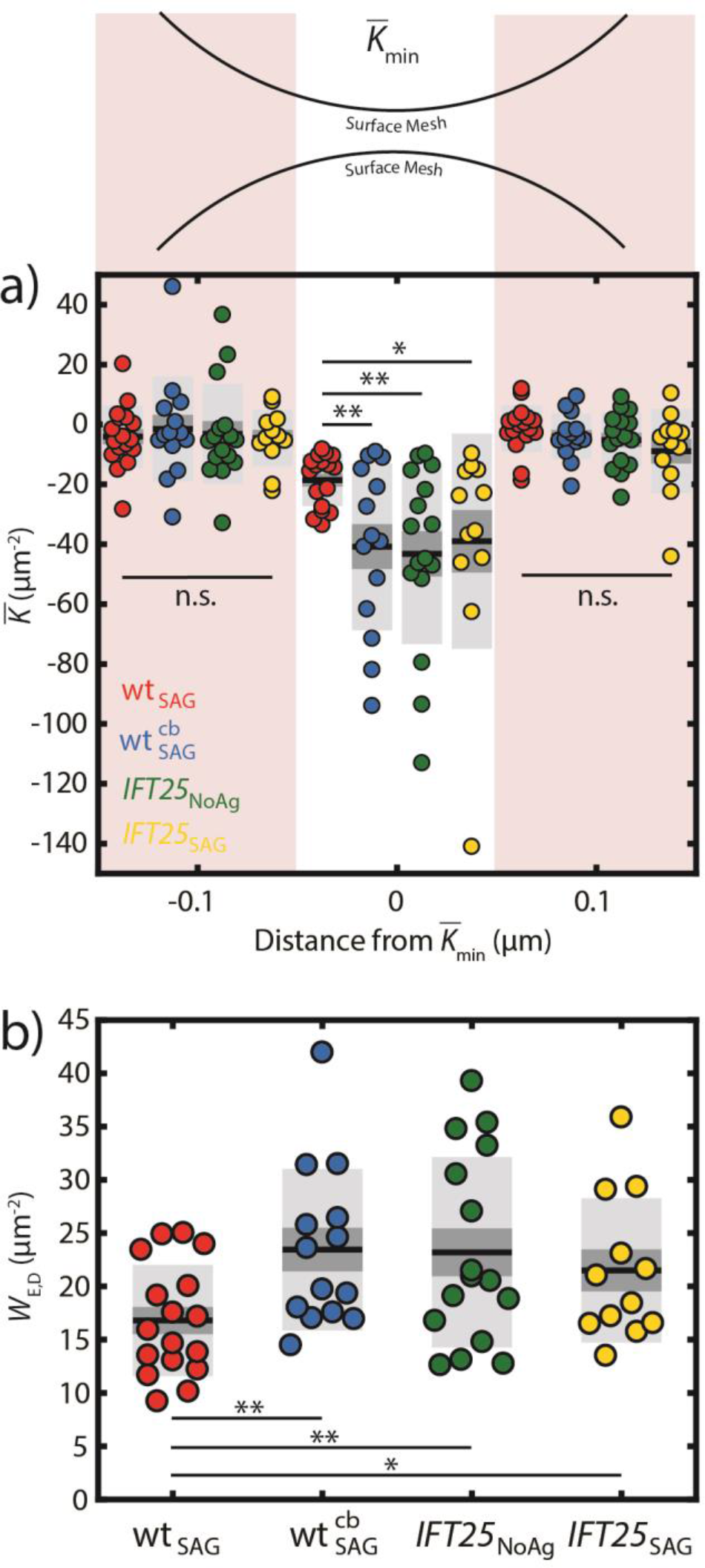
Quantitatively detecting changes in primary cilia morphology in MEF cells. When calculating the *K̅_min_* and both neighboring regions 0.1 μm for each cilium, (**a**) there is a significant decrease in *K̅_min_* found in mutant cells compared to the control wt MEF cells. (**b**) *W_E, D_* measures the overall shape of the object and is found to be significantly higher in mutant cells. Reporting mean ± S.E.M. (dark gray), and mean ± S.D. (light gray). (** *p* < 0.01, * *p* < 0.05)

Our 3D SR methods for quantifying the morphology of the primary cilium clearly reveal measurable morphological changes when the retrograde transport machinery is severely impaired in MEF cells, either by genetic knockout of *IFT25* or inhibition of dynein-2 function. Changes in ciliary membrane morphology can be subtle under certain physiological conditions, but by combining 3D SM microscopy and existing quantitative methods from differential geometry, we were able to reveal and characterize features that would have been impossible to observe using conventional microscopy techniques. Although we show *IFT25* to have at the very least an indirect effect on the membrane shape, the direct mechanism of these structural defects in the ciliary membrane is a subject of future study.

### Characterizing the membrane & axoneme simultaneously using 2-color 2D STED microscopy

Based on previous studies (63), we hypothesized the axoneme, a 9-fold symmetric microtubule doublet structure providing the central core of the primary cilium, may also be altered in our *IFT25* mutant cells. We observed both SMO and the α-tubulin component of the axonemal microtubules using primary/secondary antibody staining in fixed MEF cells. To achieve a resolution beyond the diffraction limit, we utilized STED microscopy which provided both a secondary verification of the morphological changes detailed above and also allowed us to correlate the relative distributions of SMO and the inner cytoskeleton of the same primary cilium. With a full-width half-maximum (FWHM) resolution of 50-100 *nm*, we were able to resolve the two sides of the ciliary membrane in the SMO channel and obtained an enhanced resolution of the axoneme in our STED images compared to the corresponding confocal images (Fig. S7 in the Supporting Material). In addition, we were able to measure the diameters of the membrane and the axoneme (Fig. S9 in the Supporting Material), which are consistent with previous EM studies of the primary cilium (64). We find that, as expected, the ciliary membrane diameter is larger than the axonemal diameter by ~25%, but these diameters are not significantly different when comparing *wt* to *IFT25* cells. For the *wt* MEF cells, the majority of the primary cilia had a normal phenotype, where the ciliary membrane and the axoneme were largely cylindrical in shape, although the latter was often found to stop a few hundred nanometers before it reached the tip (Fig. 6a). Although a very small proportion of *wt* cilia had a bulged membrane near the tip of the cilium, the axoneme did not extend all the way to the tip, similar to other *wt* cells (Fig S8a in the Supporting Material). Despite this, the tip still maintained a semi-hemispheric structure in 2D which is consistent with our 3D SR results. In contrast, most of the *IFT25* mutant cells imaged exhibited clear bulging near the tip of the ciliary membrane which also agrees with our 3D SR results. Furthermore, a larger proportion of these cells had antibody-labeled α-tubulin proteins spanning the entire primary cilium length and occupying the bulging ciliary tip (Fig. 6b). Although the images of the axoneme in some *IFT25* mutant cells are found to lose the normal cylindrical structure near the tip and resemble a large bulge, it is possible that this region is occupied by monomeric α-tubulin proteins (Fig S8b in the Supporting Material). Based on these observations, one may speculate that when IFT25 is absent, a larger number of ciliary proteins accumulate at the tip, leading to a bulge in the ciliary membrane. Therefore, IFT25 clearly plays some role in ensuring that the primary cilium maintains a normal shape, even though previous studies show this protein is not required for ciliogenesis.

**Figure 6:**
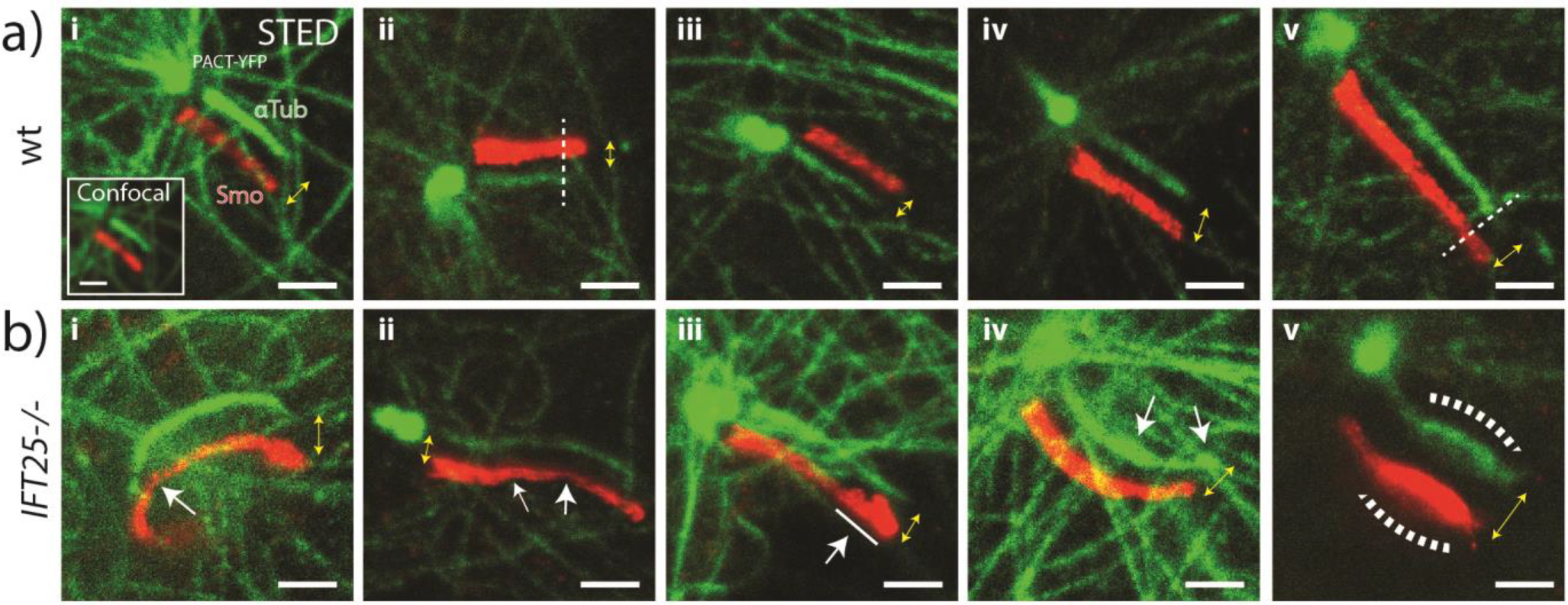
2D STED microscopy reveals structural changes in the axoneme for *IFT25* knockout cells. SNAP-SMO (Smo) and αTubulin (αTub) are stained with Atto647N and Star520SXP-labeled antibodies respectively in chemically fixed MEF cells, and are subsequently imaged using a 2-color confocal and 2-color STED microscope. Yellow double arrows indicate both magnitude shift and direction between the two red and green images. (**a**) In control wt MEF cells, (i-v) the ciliary membrane exhibits a largely cylindrical shape while the axoneme is frequently observed to not extend to the tip of the cilia. (ii) Dotted line indicates where the axoneme ends. (**b**) In *IFT25* cells, results similar to the 3D SR data are observed, where (i) kinks and (ii) narrowing of both the ciliary membrane and axoneme are pointed out in white arrows. (iii) Bulging and budding often occurs at the tip, and the axoneme appears to extend all the way to the tip and (iv) sometimes has an uneven diameter throughout the cilium. (v) Severe swelling of the ciliary membrane and the axoneme is also observed, although this is relatively rare. Scale bar = 1 μm. Image contrast within each cell line condition is identical for both channels.

## Conclusion

We have demonstrated a quantitative approach for interrogating the primary cilium morphology in mammalian cells using super resolution fluorescence microscopy of a transmembrane protein of the ciliary surface. By combining 3D SR microscopy and a meshing algorithm, we obtain high-resolution images of the ciliary membrane surface revealing a variety of nanoscale features. This high-resolution approach has revealed the existence of structural defects that were not seen in previous reports of diffraction-limited images of *IFT25* mutants in MEF cells (64); specifically our analysis indicates significant changes in cilia morphology manifested in our measurements of *H, K, W_E, D_*. We are also able to demonstrate that *wt* cells treated with ciliobrevin D exhibit similar phenotypes to the *IFT25* mutant cells. Therefore, impaired retrograde transport, either as a result of inhibiting retrograde motor dynein or due to a genetic deletion of IFT25, affects the overall shape of the ciliary membrane, primarily in form of bulging, which is indicative of a underlying mechanism related to the mobility of cargo in the cilium. We also resolved the axoneme structure within mammalian cells by implementing STED microscopy. Notably, the antibody-labeled α-tubulin proteins in the *IFT25* mutant cells are observed to span the entire primary cilium. This suggests that IFT25 plays a role in properly maintaining the structure of the primary cilium, especially near the tip, even though it is not required for ciliogenesis. Our method for characterizing the biophysical and morphological properties of the ciliary membrane can be used to study other nanoscale structures, such as bacteria or the nuclear envelope, in a similarly quantitative manner, if the surfaces can be labeled with suitable dyes for SR microscopy.

### Acknowledgements

We thank Dr. Gregory Pazour for providing us with the *IFT25* mutant cell line, Dr. Rafe Mazzeo for helpful discussions, and Dr. Pete Dahlberg for the 3D printed models. This work was supported in part by a National Science Foundation Graduate Fellowship (J.Y., C.J.C.), a Stanford Bio-X Fellowship (L.E.W.), the National Institute of General Medical Sciences through Grants No. R35-GM118067 (W.E.M.) and No. R01-GM121424 (T.S.).

### Author Contributions

J.Y., L.E.W., & W.E.M. conceived the study. L.M. created the cell lines. J.Y. prepared biological samples for imaging, performed 3D SM microscopy experiments, and analyzed all data. C.J.C. performed 2-color 2D STED experiments. J.Y., C.J.C., L.E.W., L.M., T.S., and W.E.M. wrote the manuscript.

## SUPPORTING MATERIAL

### 1. Cell Culture

Mouse embryonic fibroblast (MEF) cells are used for all our samples, which stably express Smoothened proteins with a SNAP tag (SNAP-SMO) and Pericentrin-YFP (PACT-YFP). Cell lines used: (1) **wt**: SMO−/−, SNAP-SMO, PACT-YFP; (2) ***IFT25***: IFT25−/−, SNAP-SMO, PACT-YFP ^19^. Cells are cultured in DMEM/High Glucose (Hyclone, SH30243.01) with 10% Fetal Bovine Serum (FBS) (Hyclone, SH30070.03), in 25 *cm*^2^ surface area cell culture flasks (Falcon) maintained in an incubator at 37°C and 5% CO_2_ (ThermoFisher, Heracell 160i). When cells reach 80-90% confluency, cells are detached from the surface using 0.25% Trypsin (Hyclone, SH30042.01) and are pipetted into a 4-well chambered borosilicate coverglass (Fisher Scientific, 12-565-401). Immediately after they are plated, 200 *nm* diameter red fluorescent beads, ex: 580 *nm*, em: 605 *nm* (Invitrogen, F8801) are added to the sample. Cells are cultured in 10% FBS media for 24-48 *hrs* before they are serum starved in 0.5% FBS media for an additional 20-24 *hrs*. (1) **wt** cells are treated with 10 *uM* Smoothened Agonist (SAG) for 4 *hr* at the 20-*hr* mark of serum starvation. A separate chamber containing **wt** cells are treated with 10 *uM* ciliobrevin for 1 *hr* before the 24-*hr* mark. (2) ***IFT25*** cells are either (a) treated with SAG or (b) none at all (NoAg).

### 2. Sample Preparation (3D SM Microscopy)

Cell samples are first labeled with BG-Alexa647 (NEB, S9136S) at 3 *uM* concentration for 20 *min* at 37°C and 5% CO_2_. Each chamber of cells is washed with 0.5% FBS three times at 5-*min* intervals. Cells are fixed using 4% paraformaldehyde (PFA) (Alfa Aesar, 43368) for 15 *min* and then treated with a quenching solution of 10 *mM* NH4Cl for 10 *min*, both steps at 25°C. The samples are washed with PBS, pH 7.4 (1X) (Gibco, 1789842) at 5-*min* intervals and then stored at 4°C up to 1-week before being discarded.

### 3. 3D SM Microscopy Setup

Experiments are performed on a customized inverted microscope (Olympus, IX71) where the sample is mounted on a piezo-electric stage (PI-Nano) and is in contact with an oil-immersion objective (Olympus, 100x, 1.4 NA, UPLANSAPO) applied with a small drop of oil (Immersol, 12-624-66A) before mounting. New imaging buffer is added for each primary cilium imaged (1-2 *hrs*), which consists of glucose oxidase (Sigma-Aldrich, G2133), catalase from bovine serum (Sigma-Aldrich, C100), 100 mM Tris-HCl, pH 8.0 (ThermoFisher Scientific, 15568025), 10% (w/v) glucose solution (Sigma-Aldrich, 49139), 140 *mM* beta-mercaptoethanol (Sigma-Aldrich, M6250), and H_2_O (Nanopure) 45. We locate one primary cilium and image SNAP-SMO-Alexa647 and PACT-YFP using the 641 *nm* (Coherent Cube, 100 *mW*) and 514 *nm* (Coherent Sapphire, 50 *mW*) laser, respectively. Fluorescence (emission) is collected through the objective, a dichroic filter (Semrock, FF425/532/656-Di01), and two bandpass filters (Chroma, 680-60; Chroma, 655LP). When the primary cilium is placed at the center of the field-of-view (FOV) and is in focus, the double-helix (DH) phase mask is carefully placed at the Fourier Plane (FP) in our *4f* system, with our lenses each having a focal length of *f* = 90 mm. We increase the intensity of the 641 *nm* laser (1-5 *kW/cm*^2^) and allow the fluorescent dye to bleach down to the single-molecule regime. Over the next hour, we gradually increase the 405 *nm* (Obis, 100 *mW*) laser intensity until either 1 *hr* has passed or single-molecule blinking becomes extremely sparse. Red fluorescent beads are also imaged simultaneously several microns away from the PC. Detected fluorescence is recorded using a silicon EMCCD camera (Andor Xion, DU-897U-CS0-#BV) at a speed of 14-20 frames/second (50-70 *ms*/frame) with an electron-multiplying gain of 200.

**Figure S1:**
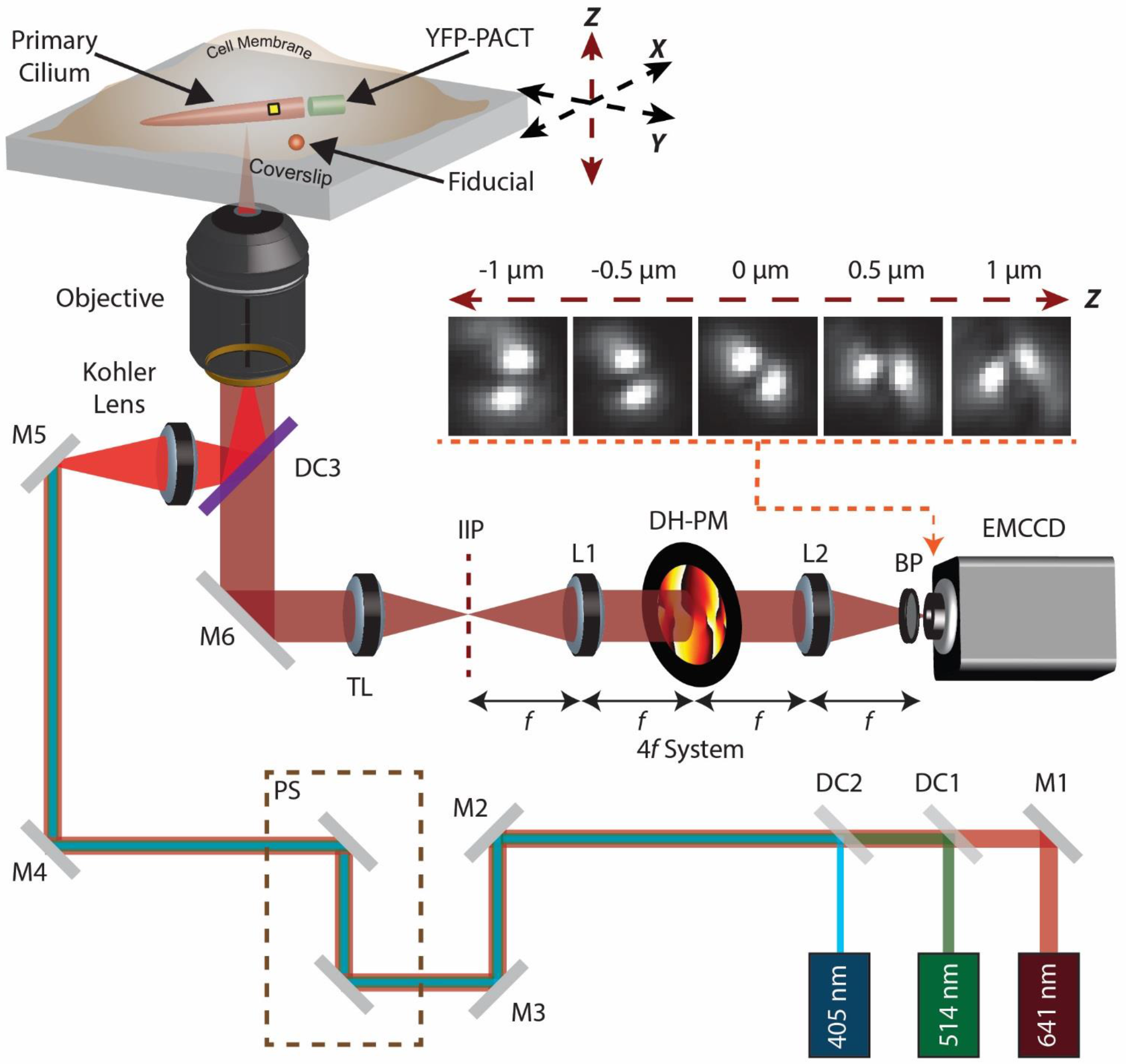
3D SR Microscopy setup with the Double-Helix Phase Mask (DH-PM). Lasers go through a series of mirrors (M), dichroic mirrors (DC), and a periscope (PS), before it passes through the Kohler lens and the objective. The sample consists of a primary cilium that is sandwiched between the attached cell and the coverslip surface, and a nearby stationary fiducial. Emission is collected with the objective which then goes through the tube lens (TL) and rather than placing the camera at the intermediate image plane (IIP), the *4f* system is implemented, where the double-helix phase-mask (DH-PM) is placed equidistant between two *4f* lenses (L). A bandpass filter (BP) is placed right in front of the camera (EMCCD) which is used to detect the emitted fluorescence.

### 4. Z Calibration with Fiducials

DHPSF calibrations are done using two different beads: (1) 625/645 *nm*, which is for the SNAP-SMO, and (2) 580/605 *nm*, which is for fiducials in our sample, correcting for sample drift. Calibration imaging samples are prepared by spin-coating the beads in 1% polyvinyl alcohol (PVA) onto a glass coverslip. Samples are then mounted onto the 3D SR microscopy imaging setup with the DH-PM placed at the FP. Using our piezo-electric stage, we scan over a 3 *μm* range along the *z* axis with a 50 *nm* step-size with 30 frames measured at each *z*-height. We perform a forward and backward scan to account for hysteresis during the *z*-scan. This calibration step also produces template images of the DH-PSF which are used for the identification of single-molecule signals during post-processing of the raw data. All imaging is done at 25°C.

### 5. 3D Localization of SNAP-SMO Molecules

Using the *easyDHPSF* ^41^ MATLAB program, a *z*-axis calibration over a 3-*μm* range is obtained via a 2D Double-Gaussian fit, which provides us with *xy* positions, width, amplitudes, and offset levels of each lobe of the fluorescent bead. Collected raw data is used to calculate the phase-correlation to locate single-molecules in the FOV and is ultimately fitted using the same model. An array of 3D positions is obtained, including information on photon counts, background counts, and other parameters relevant to our fits.

### 6. In-situ Localization Precision Calculations

For one 3D localization dataset from one primary cilium, we choose one localization of interest and pool in any other localizations that are both (1) within the next 10 frames and (2) within a ~74 nm 3D radius 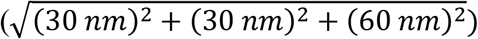. Each cluster of localizations has a minimum of 5 frames that meet our criteria. For calculating the localization precision, we calculate the standard deviation for each set of x, y, and z positions. Average signal and background photons is calculated by evaluating the mean over all localizations within each cluster.

### 7. 2D Surface Meshing

Scatterplots are prepared in the following way. We first select a few points near the base of the cilium, perform an elliptic fit, which also defines a plane of rotation, and then produce a direct copy of the original scatter plot which is rotated 180°. This step is important for creating a closed surface and to minimize artifacts near the base when performing our final curvature analysis. This new scatterplot is then used to extract out the three-dimensional surface using MeshLab. This program is used to first calculate the normal vectors of each point using a range of 30-200 neighboring points, which varies depending on how many localizations there are in one point cloud. Then, a Poisson Surface Reconstruction algorithm is applied to our processed data in order to create a triangulated surface (Octree Depth = 13; Solver Divide = 6; Samples per Node = 5; Surface Offsetting = None). Gaussian curvature is calculated to highlight areas of both positive and negative curvature. This data is then exported in the standard ‘*.ply’ file format which is used to plot our 3D mesh and further curvature analysis.

**Figure S2:**
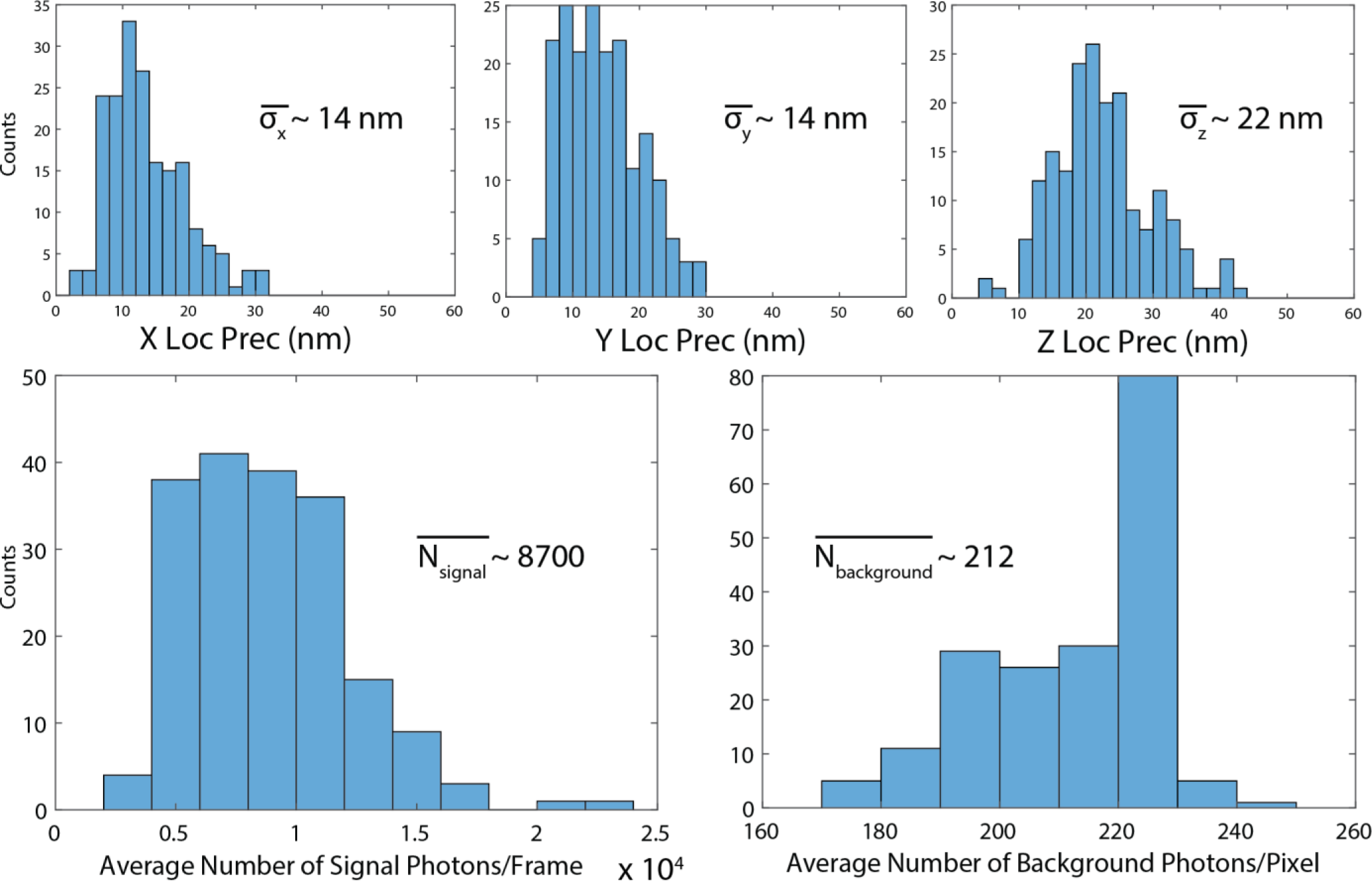
Localization precision measurements for x, y, and z with average number of signal and background photons per frame.

### 8. 3D-Printed Meshes

MeshLab creates the .STL file format which lists the x, y, z coordinates of the vertices on the mesh. An Ultimaker-2 printer using PLA filament and an infill setting of 25% printed the model. Printing was performed at the Atherton public library in San Mateo County.

**Figure S3:**
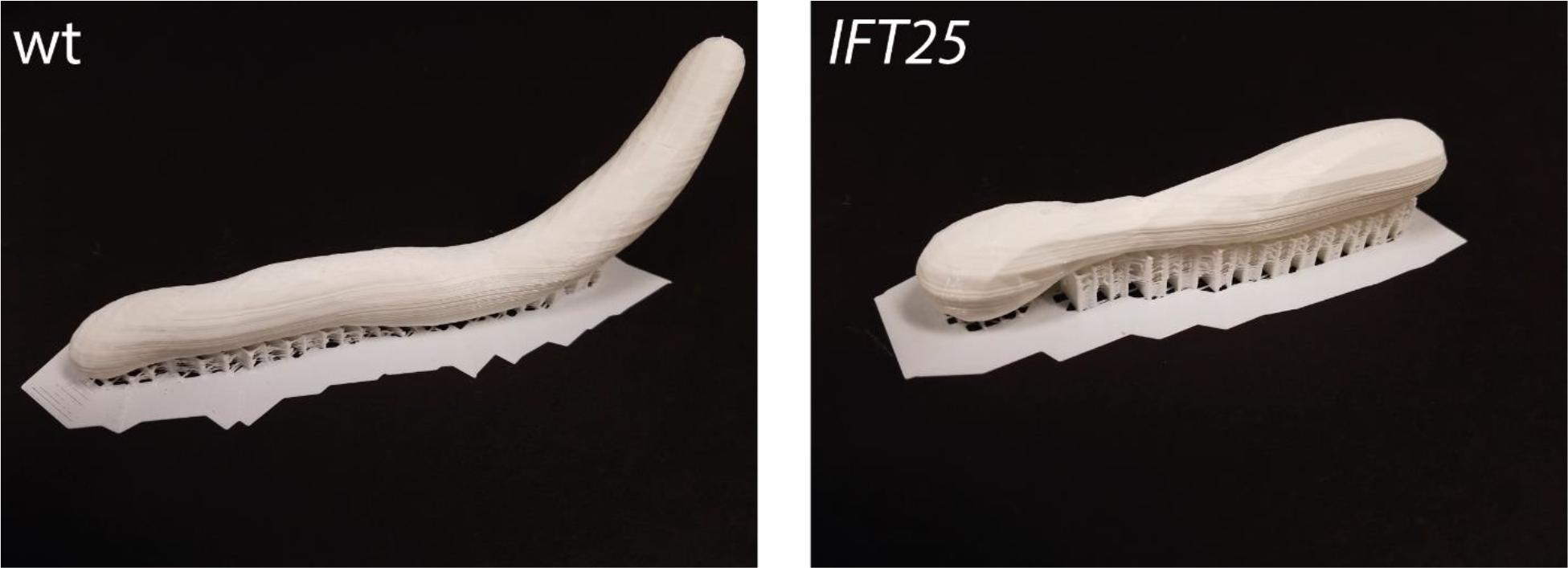
3D printed cilia.

### 9. Curvature Analysis

**Figure S4:**
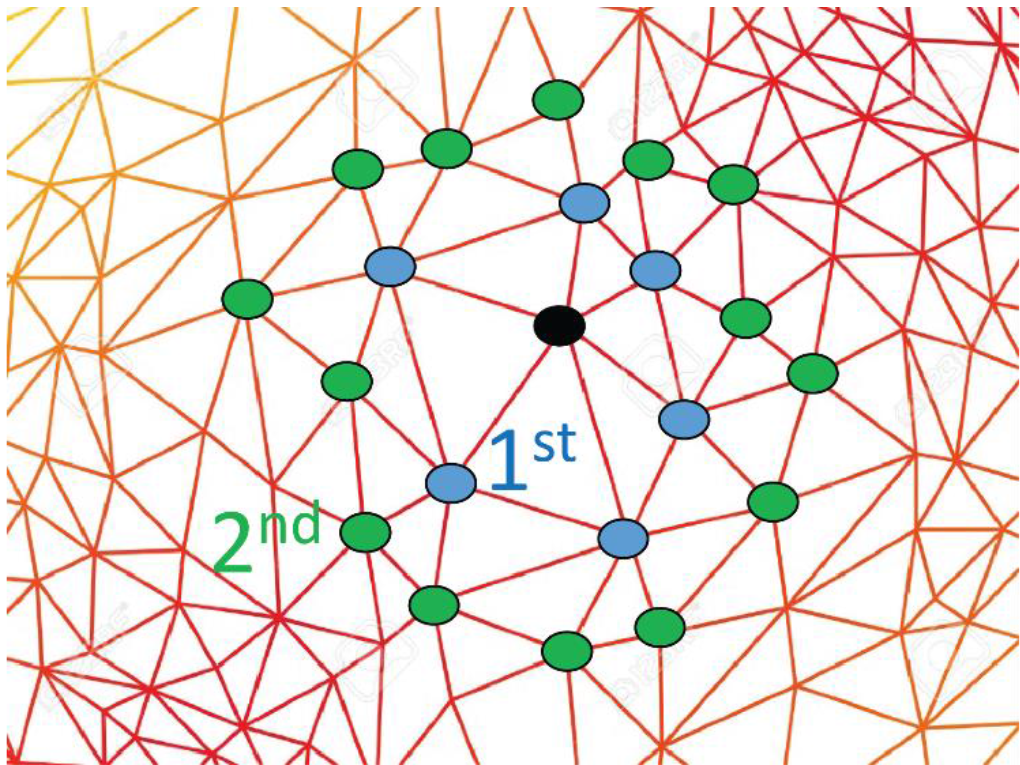
Illustration of mesh with 1^st^ and 2^nd^ order neighboring points to the black point.

We first choose a vertex along the mesh and search for its 1^st^ & 2^nd^ order nearest neighbors, which all make up a “patch” of triangles. Within each patch, we calculate the normal vector for every triangle and the angle at the vertex of interest. We determine the overall weighted normal vector of the patch and is used to rotate all points such that this vector is oriented along the *z*-axis. We then fit a surface to our points of the following form:

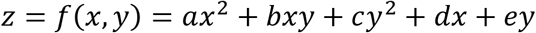

If we further define *p* = [*a b c d e*], *A* = [*x*^2^ *xy y*^2^ *x y*], we solve for *p* by evaluating the following:

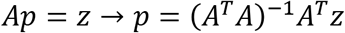

Using the coefficients in *p*, we then calculate the Mean Curvature (H) and Gaussian Curvature (K) using the following expressions ^46^:

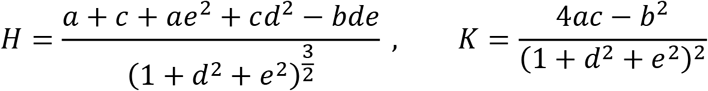

This was done for every vertex and are represented as heat maps, interpolated along the mesh. We calculate the Willmore Energy by calculating the following

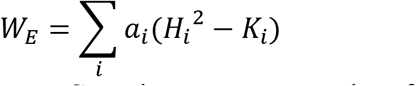

where *H_i_*, *K_i_*, and *a_i_* are the mean curvature, Gaussian curvature, and surface area of the *i^th^* triangle of the mesh respectively.

### 10. Ciliary Length and Surface Area

**Figure S5:**
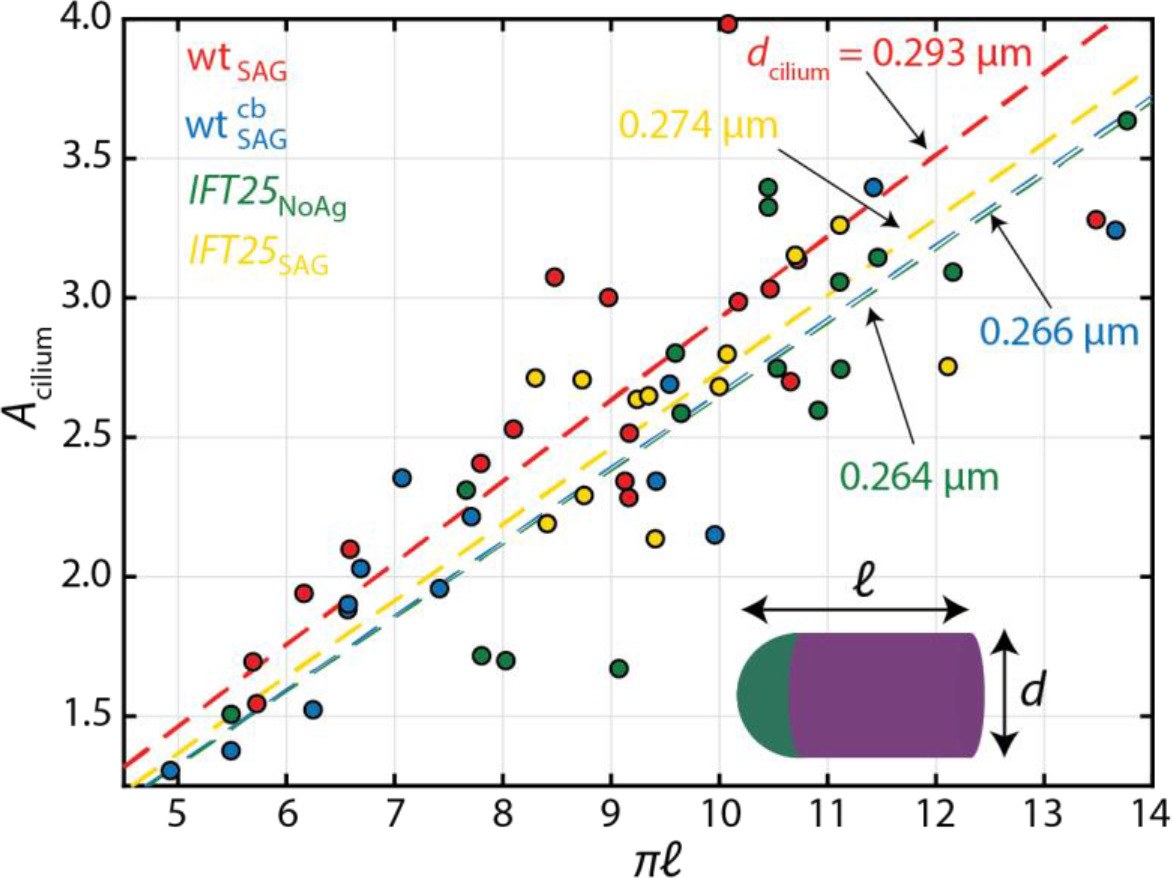
Ciliary length and surface area for each condition, cell by cell. The total surface area of the cilium, *A_cilium_*, is calculated by adding up the areas of all the triangles of the ciliary mesh. The ciliary length, *l*, is calculated as the length of the ciliary axis from the base to the tip. The population diameter, *d_cilium_*, is estimated by the output slope of a linear regression performed for each data set. Here we are making an assumption that the cilium takes on the approximate shape of a cylinder with a hemispheric cap, as shown in the figure inset on the bottom-right corner.

For values of the diameter for each cell, *d*, in Table 1, the following derivation and calculation was utilized:

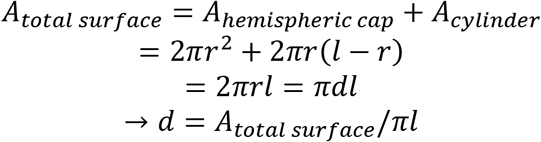

### 11. Sample Preparation (2D STED)

Cell samples are fixed with 4% PFA for 15 *min* at 25°C, washed with 1x PBS, then immersed in a blocking solution, consisting of 1% TritonX-100, Normal Donkey Serum (Jackson ImmunoResearch, 017-000-121), and 1x PBS, for 30 min at 25°C. We stain our samples with primary antibody for 1 *hr*, consisting of anti Smo-C and anti AlphaTubulin (Sigma-Aldrich, T6199) in blocking solution. Samples are then washed 3x with 1x PBS at 5-*min* intervals, then stain our samples with secondary antibody for 1 *hr*, consisting of goat anti-rabbit atto647N (Active Motif, 15048) and goat anti-mouse Star520SXP (Abberior, 2-0002-009-9) in blocking solution. Samples are then washed 3x with 1x PBS at 5-*min* intervals then are either immediately imaged or are stored at 4°C for up to 1 week

### 12. 2-color 2D STED Microscopy

**Figure S6:**
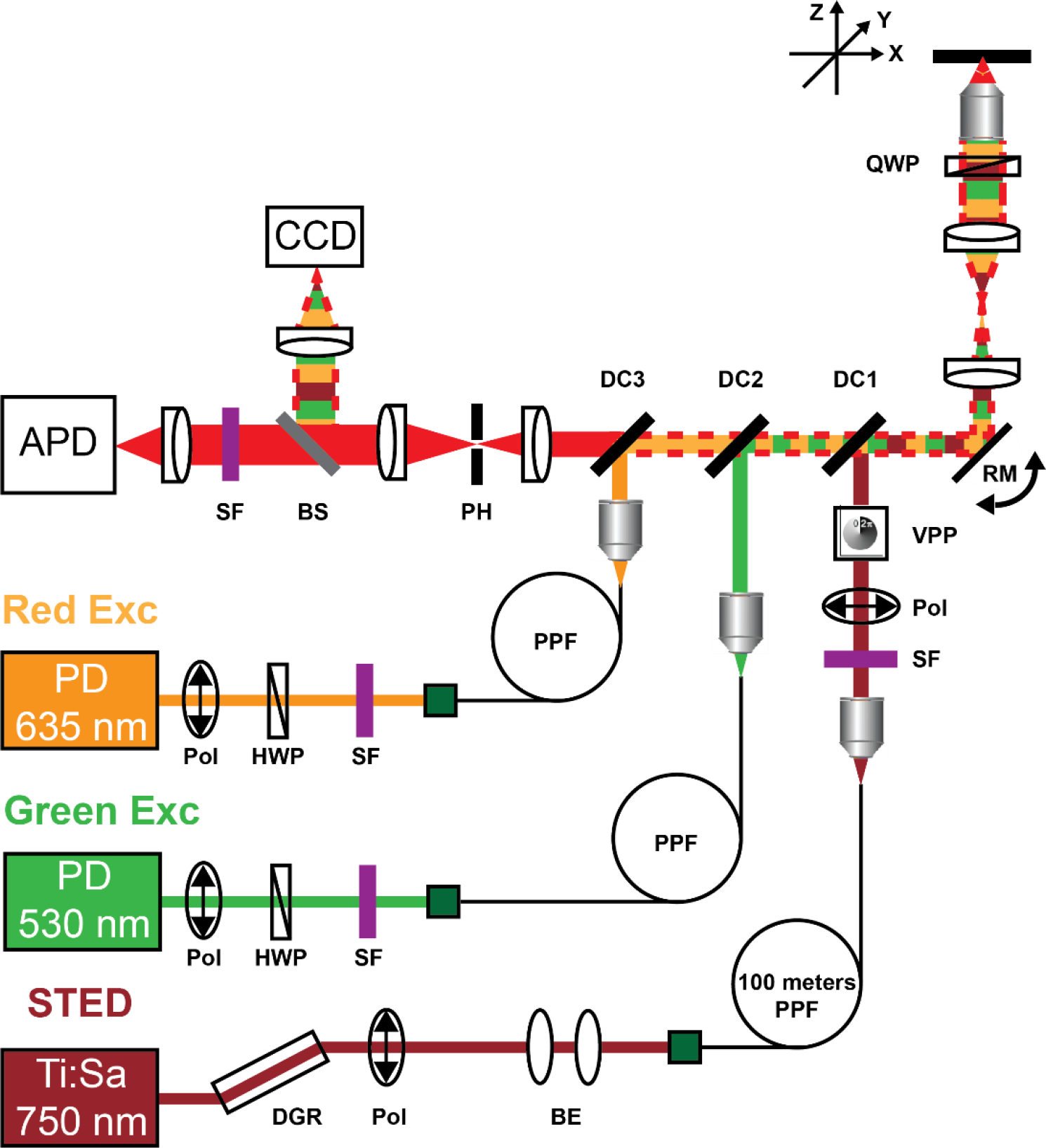
2D STED Microscopy Schematic. Lasers are sent through a cleanup polarizer (Pol) and spectral filter (SF). The two excitation lasers’ polarization are rotated by half-wave plates (HWP) prior to coupling into a polarization preserving fiber (PPF). The depletion laser pulse is temporally stretched by a dispersive glass rod (DGR) before the beam is expanded by a beam expander (BE) and is further temporally stretched by 100 meters of PPF. The fiber outputs are collimated using objective lenses. The depletion beam is sent through another Pol and the donut shape is imparted by a vortex phase plate (VPP). All three beams are coupled by a series of dichroics (DC), and are scanned by a resonant mirror (RM), made conjugate to the back focal plane of the imaging objective by a telescope lens pair. The beams are made circularly polarized by a quarter-wave plate (QWP). Emission is collected with the imaging objective, is de-scanned by the same RM, and passes through the DCs. It passes through a confocal pinhole (PH), is filtered by a SF, which changes depending on the color channel, and is detected on an avalanche photodiode (APD). A beam splitter (BS) is used to image a small fraction of reflected/emitted fluorescence onto a camera (CCD) for alignment purposes.

STED images were collected on a bespoke 2-color fast scanning STED microscope (Figure S6). The 750 *nm* depletion laser is provided by a titanium-sapphire mode-locked oscillator operating at 80 MHz (Mira 900D, Coherent). The pulses are dispersed to ~200 *ps* in duration using 30 *cm* of SF2 glass, 10 *cm* of SF6 glass, and 100 *m* of polarization maintaining optical fiber (OZ Optics). The pulses are spectrally filtered (FF01-715/LP, Semrock) and the donut shape is created using a vortex phase-plate (RPC Photonics). The excitation pulses are provided by 530 *nm* and 635 *nm* pulsed diode lasers (LDH-P-FA-530B & LDH-P-C-635B, PicoQuant) that are electronically triggered to arrive prior to the depletion pulses. The excitation beams are spatially filtered by polarization maintaining fibers (Thorlabs), and combined using a 532 *nm* longpass and 514/640 *nm* notch dichroic (ZT532RDC & ZT514/640RPC, Chroma). The excitation beams are combined with the depletion beam by a 5 *mm* thick 710 *nm* shortpass dichroic (Z710SPRDC, Chroma). The beams enter the back-port of a Nikon TE300 inverted microscope and are converted to circularly polarized light using a quarter-wave plate (767 *nm* zero-order, Tower Optical). The light is focused through an oil immersion objective (Plan Fluor 100x/1.3 NA, Nikon). At the sample plane the green and red excitation beams have an average power of 40-60 *kW*/*cm*^2^ and 50-80 *kW*/*cm*^2^, respectively. The depletion beam has an average power of 120-130 *MW*/*cm*^2^. The fast axis is scanned using a 7.5 kHz resonant mirror (Electro-Optical Products) that is imaged onto the back focal plane of the objective using a Keplarian telescope. The slow axis is scanned using a piezo stage (PD1375, Mad City Labs). Fluorescence is detected through the same objective, is de-scanned by the same resonant mirror, and passes through the previously mentioned dichroics. An aperture corresponding to a ~0.7 AU and 0.8 AU (red and green channels respectively). The fluorescence is then spectrally filtered by a 715 *nm* shortpass (FF01-715/SP, Semrock) followed by either a 635 *nm* longpass (BLP01-635R, Semrock) or a bandpass (ET585/65m, Chroma) for the red and green channels respectively. Fluorescence is detected on a Si APD detector (SPCM-ARQH-13, Perkin Elmer). Scan control and image acquisition use a custom LabVIEW algorithm running on an FPGA (PCIe-7842R, National Instruments) and host computer. STED images have a pixel size of 20 *nm*. Images in the red channel are obtained as a single frame with 1000 resonant mirror scans per line or an average pixel dwell time of ~0.1 *ms*/pixel. Images in the green channel are obtained as 2-3 frames with 200 resonant mirror scans per line for each frame or an average pixel dwell time of ~20 *μs*/pixel/frame.

### 13. 2D STED Resolution

**Figure S7:**
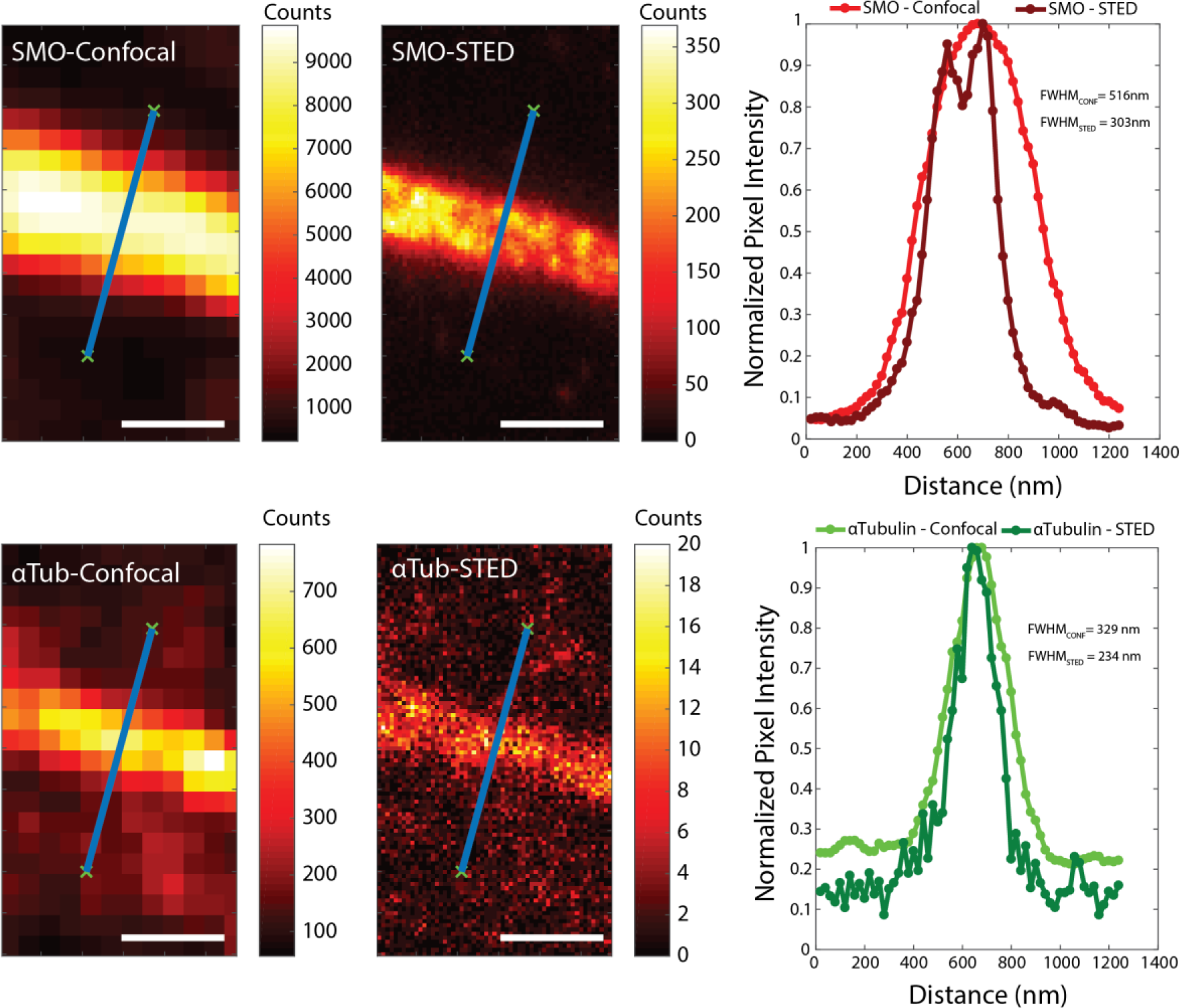
Comparing confocal and STED image line profiles for one primary cilium in both color channels for SMO (upper) and α-tubulin (lower). Line profiles were generated by extracting signal counts that reside in 20 nm bins underneath the drawn line for both channels.

### 14. Additional 2D STED Images of Unusual Primary Cilia

**Figure S8:**
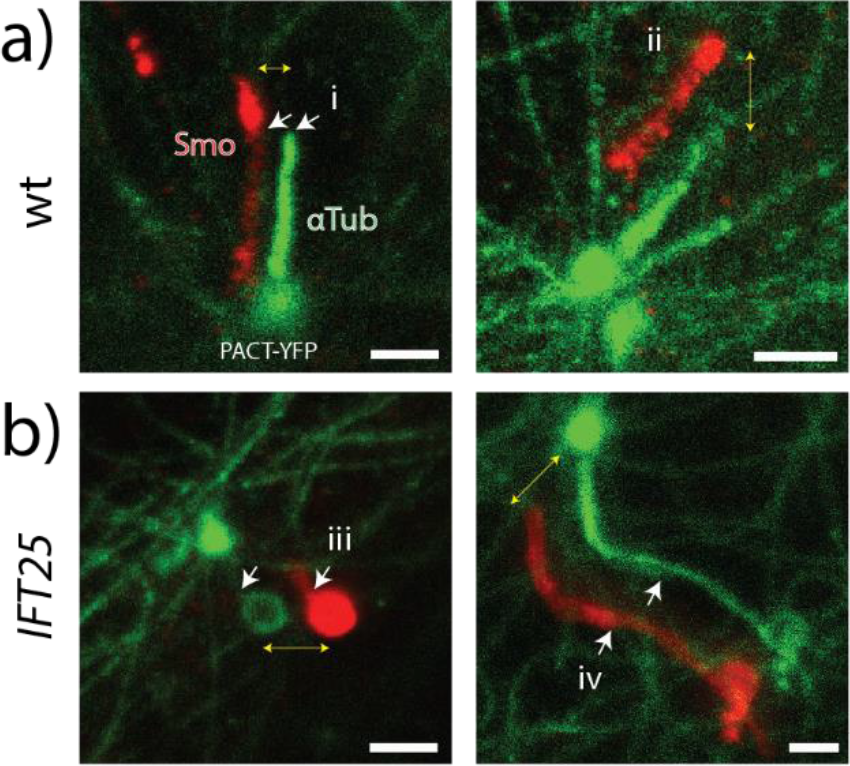
Additional 2D STED images of unusual primary cilia. (a) Control MEF cells also exhibit (i, ii) bulging at the tip at times, where the axoneme does not extend to the tip. (b) Some cases of *IFT25* mutant cells produce oddly-formed cilia which are (iii) very short with a circular tip or (iv) very long with a narrowing along the shaft of the cilium with a bulging tip

### 15. Membrane/Axoneme Diameter Calculations

**Figure S9:**
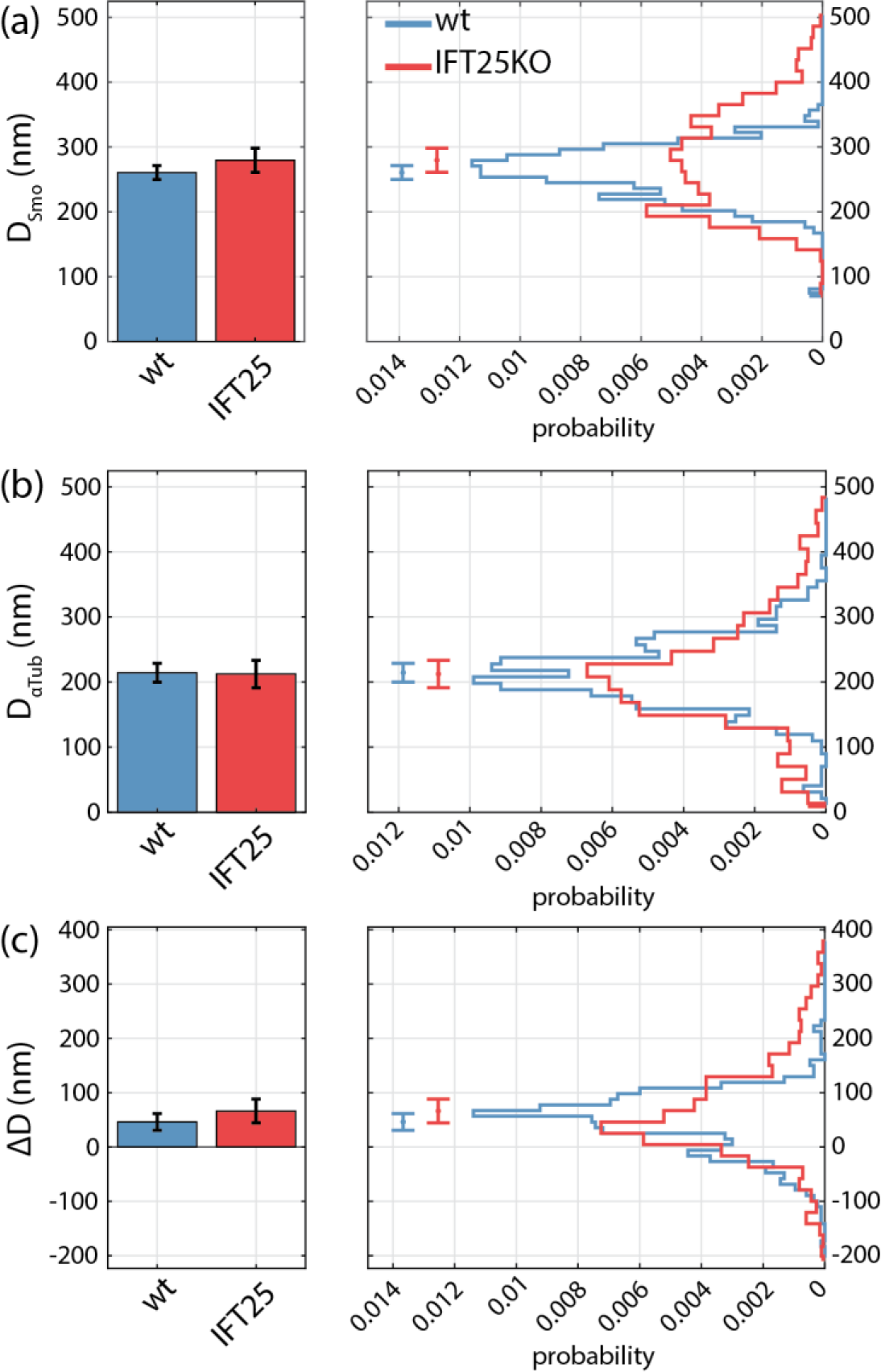
Distributions of diameter measurements using 2D STED. Histograms and bar plots of (a) SNAP-SMO (*D_Smo_*), (b) α-tubulin (*D_αTub_*), and (c) difference (*D_Smo_* − *D_αTub_*). There is ~18% and 24% difference between SNAP-SMO and α-tubulin for *wt* and *IFT25* mutant cells, respectively. Reporting mean ± S.E.M (*N_wt, cilia_* = 12, *N_wt, profiles_* = 801, *N*_*IFT*25, *cilia*_ = 15, *N*_*IFT*25, *profiles*_ = 993).

2D STED SNAP-SMO images are used to determine the ciliary axis, which is done by performing a spline fit to a set of user-selected input points chosen along the center of the cilium by eye. The intensity line profile, 600 nm long with a bin width of 20 nm perpendicular to the ciliary axis is determined at 50-nm long steps along the ciliary axis for both channels, resulting in a representation of the lateral extent of the cilium at each position. A 1D double-Gaussian fit to each profile is performed of the following form:

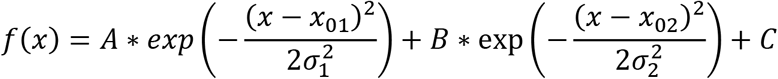

where A, B are the amplitudes, C is the constant background, *x*_01_, *x*_02_ are the centroids, *σ*_1_, *σ*_2_ are the standard deviations. The full-width half-maximum (FWHM) is calculated for each profile, *FWHM* = *x*1_*f*(*x*1)=1/2_ − *x*2_*f*(*x*2)=1/2_, which we define in this study as the diameter for the SMO-determined membrane and the axoneme.

### 16. Supplementary Movies

**Movie 1: Control wt MEF primary cilium 3D mesh**

Camera flythrough, which starts above the mesh, then follows along the ciliary axis, and finally pans away. This is the same mesh as shown in Figure 2a. White bounding box dimensions, 2750 nm x 1500 nm x 1500 nm; video speed, 30 frames/sec. The HSV color scaling represents z in the image over a range from −175.8 to 1194.5 nm.

**Movie 2: *IFT25* mutant MEF primary cilium 3D mesh**

Camera flythrough, which starts above the mesh, then follows along the ciliary axis, and finally pans away. This is the same mesh as shown in Figure 2b. White bounding box dimensions, 3700 nm x 900 nm x 600 nm; video speed, 30 frames/sec. The HSV color scaling represents z in the image over a range from −196.4 to 227.1 nm.

